# Epigenetic regulation of visual system remodeling during flatfish metamorphosis: DNA methylation dynamics in ocular migration and visual adaptation

**DOI:** 10.1101/2025.02.19.638519

**Authors:** Laura Guerrero-Peña, Paula Suarez-Bregua, Nuria Sánchez-Baizán, Francesc Piferrer, Juan J. Tena, Josep Rotllant

## Abstract

Flatfish metamorphosis is characterized by extensive tissue remodeling, associated with a transition from pelagic to benthic lifestyle, being the migration of one eye the most dramatic change. Epigenetic mechanisms exert a pivotal role in developmental programs. This study investigates the DNA methylation profiles of migrating and non-migrating eyes using reduced-representation bisulfite sequencing (RRBS) during three developmental stages of turbot: pre-metamorphosis, climax and post-metamorphosis. Over 31% of all identified regions were hypermethylated during climax stage in both eyes, coinciding with elevated expression of the *dnmt3a* gene, responsible for de novo methylation. Additionally, transcription factors crucial for retinal ganglion cells (RGCs) development, including the *eomesa* and *tbr1b*, exhibited differential methylation and expression between the migrating and non-migrating eye during the climax phase. These findings underscore the significance of DNA methylation in the intricate remodeling of the visual system during turbot metamorphosis, particularly regarding RGC-mediated ocular migration and the transmission of visual signals.

## Introduction

Epigenetic modifications, including DNA methylation, histone modification, and non-coding RNAs, play a pivotal role in regulating gene expression across diverse biological processes. In vertebrates, DNA methylation, typically occurring in a cytosine-guanine residues (CpG) context, can lead to gene repression or activation. This dynamic process is essential for tissue differentiation, growth and development, and for adaptation to environmental changes, ensuring the successful progression of developmental programmes (***Meehan and Stancheva, 2001; Jones et al., 1998***).

One of the most fascinating examples of developmental transformation in vertebrates is the metamorphosis observed in flatfishes (order Pleuronectiformes). This process entails a dramatic reorganization of the body plan, characterized by asymmetrical morphological changes, where one eye es relocated from one side of the head to the other. This distinctive adaptation is vital for the benthic existence of adult flatfish, facilitating their capacity to evade predators on the ocean floor (***Friedman, 2008***). The transition from a pelagic to a benthic lifestyle also requires adaptations to varying light conditions, entailing a substantial reorganization of the retina’s photoreceptors during metamorphosis (***Bolstad and Novales Flamarique, 2022; Evans and Fernald, 1993; Hoke et al., 2006***). Both eyes are exposed to reduced light conditions, resulting in no discernible developmental differences in the ocular layers between themn (***Villegas et al., 1997***), and some studies have reported no differences in the organization of visual projections between the fixed eye and the migrating eye (***Medina et al., 1993***). However, there are distinctive characteristics associated with both eyes as one migrates to the other side of the body. For example, evidence indicates that the optic nerve in the migrating eye is significantly shorter than in the other eye (***Murray, 1974***), and the optic tectum, which receives projections from the migrating eye during metamorphosis, appears to be smaller (***Briñón et al., 1993***). This may indicate a reduction in the functionality of the migrating eye during the metamorphic remodeling process (***Briñón et al., 1993***).

Flatfish metamorphosis is a complex process that is governed by tightly regulated, tissue-specific gene expression, which is known to be controlled by the thyroid hormone (TH) (***de Jesus et al., 1993; Gomes et al., 2015; Shao et al., 2017***). A correlation has been identified between the physiological increase in triiodothyronine (T3) during metamorphosis and alterations in chromatin structure (***Cheng et al., 2010***), which facilitates the interaction of transcription factors with novel genomic regions. Moreover, TH regulates de novo methylation at the climax of metamorphosis via DNA methyltransferase DNMT3 (***Kyono et al., 2016***) in other vertebrates, which is crucial for DNA methylation pattern establishment during early development (***Rottach et al., 2009***). These findings underscore the significance of epigenetic mechanisms in metamorphic remodeling and the modulation of gene expression by DNA methylation during this process. Furthermore, DNA methylation is also known to play a pivotal role in the development of retinal ganglion cells, which form the eye-to-brain communication pathway and whose axons constitute the optic nerve, thereby emphasizing its importance in ocular development and function (***Tai et al., 2023***).

Although the significance of DNA methylation in developmental processes is well-documented (***Bogdanović et al., 2017, 2011***), the methylation landscape of the migrating and non-migrating eyes during flatfish metamorphosis and its potential relationship with gene expression remains largely unexplored. The objective of this study was to elucidate the methylation profiles of both eyes during critical stages of flatfish metamorphosis, namely the pre-metamorphic, climax of metamorphosis and post-metamorphic stages. The analysis of the methylome at the genome-wide level revealed that similar methylation patterns were identified among the stages in both eyes that have adapted to a benthic lifestyle. A greater number of hypermethylated regions were observed in differentially methylated regions (DMRs) at the climax stage, which coincides with a peak in *dnmt3a* gene expression. Additionally, we identified differential methylation resulting in differential expression between the migrating and non-migrating eye of the transcription factors *eomesa* and *tbr1b*, which are essential for retinal ganglion cell development. Our findings provide a deeper understanding of the epigenetic regulation that underpins one of the most extraordinary examples of vertebrate metamorphosis and provide the basis for further research into the adaptive evolution of flatfish.

## Results

### Differentially methylated stage-specific regions

Methylation profiles of turbot migrating and non-migrating eye were examined throughout metamorphosis, specifically at the pre-metamorphic, the climax and the post-metamorphic stages, and considering both the migrating and non-migrating eye. The mean number of raw single reads per sample for the non-migrating eye was 22.3 million, and 20.9 million for the migrating eye samples. Most clean adapter sequences (over 90%) were successfully aligned (mean of 90.54% alignment rate). A total of 38,226 and 12,616 differentially methylated regions (DMRs) were identified across stages in the migrating and non-migrating eyes, respectively. The greatest discrepancy in the number of DMRs between stages was observed between the premetamorphic and post-metamorphic stages (20,203 and 7,666 DMRs in the migrating and non-migrating eye, respectively). The median methylation difference between the identified DMRs was greater than 25% in both eyes, with a median DMR size of 1,622 bp in the migrating eye and 926.5 bp in the non-migrating eye.

### DNA methylation-mediated visual adaptation to the benthic environment

To gain insight into adaptation of turbot’s eyes to benthic environmental conditions and lifestyle established during metamorphosis, we conducted a comparative analysis to identify the sets of DMRs that exhibit a similar methylation profile across the three main stages of the metamorphic process in both eyes ***Figure 1***a. A considerable number of uniquely hypermethylated regions were identified at the climax stage, in comparison to other stages, particularly in the migrating eye ***Figure 1***a. More specifically, 15,595 distinct hypermethylated regions (with a total size of 29.1 Mb) were identified in the migrating eye. In contrast, 3,312 unique hypermethylated regions (total size of 4.47 Mb) were located in the non-migrating eye. However, the number of hypomethylated regions declined sharply during the climax stage in both eyes. A total of 3,280 hypomethylated specific regions to the climax stage were identified, exhibiting a profile comparable to that observed in the pre-metamorphic or post-metamorphic stages in both eyes. In order to gain insight of the biological processes that contribute to the enrichment of regions with a comparable methylation profile in both eyes, we identified the nearest gene and examined the associated enriched GO terms. We observed the presence of GO terms that were exclusive to these hypermethylated regions, such as axonal guidance of retinal ganglion cells and the establishment of the receptor-alpha signaling pathway of the apical/basal polarity of epithelial cells, in contrast to those that were exclusive to hypomethylated regions, such as regulation of platelet–derived growth factor receptor–alpha signaling pathway or swimming behavior ***Figure 1***b. It is noteworthy that only three of these regions exhibited hypermethylation exclusively during the climax phase, compared to the pre-metamorphic and post-metamorphic phases, in both eyes. These regions colocalized with four different transcripts, which correspond to long non-coding RNAs (lncRNAs) (ENSSMAT00000066880, ENSSMAT00000060550, ENSSMAT00000055262, ENSSMAT00000046824) ***Figure 1***c. The linc2funtion software revealed that the targeted lncRNAs could interact and form complexes with HNRNPA1, NONO or SFRS1 proteins ***Figure 1***d. Given the increased hypermethylation in the identified DMRs, particularly at the climax stage of metamorphosis, we proceeded to verify the expression profile of DNA methyltransferases genes that promote de novo methylation (*dnmt3a*) and maintenance of methylation (*dnmt1*) ***Figure 2***a. The expression of *dnmt3a* gene peaked during the climax in the migrating eye, while the *dnmt1* gene exhibited reduced its expression, with a nearly identical profile observed in both eyes ***Figure 2***b and c.

**Figure 1.**
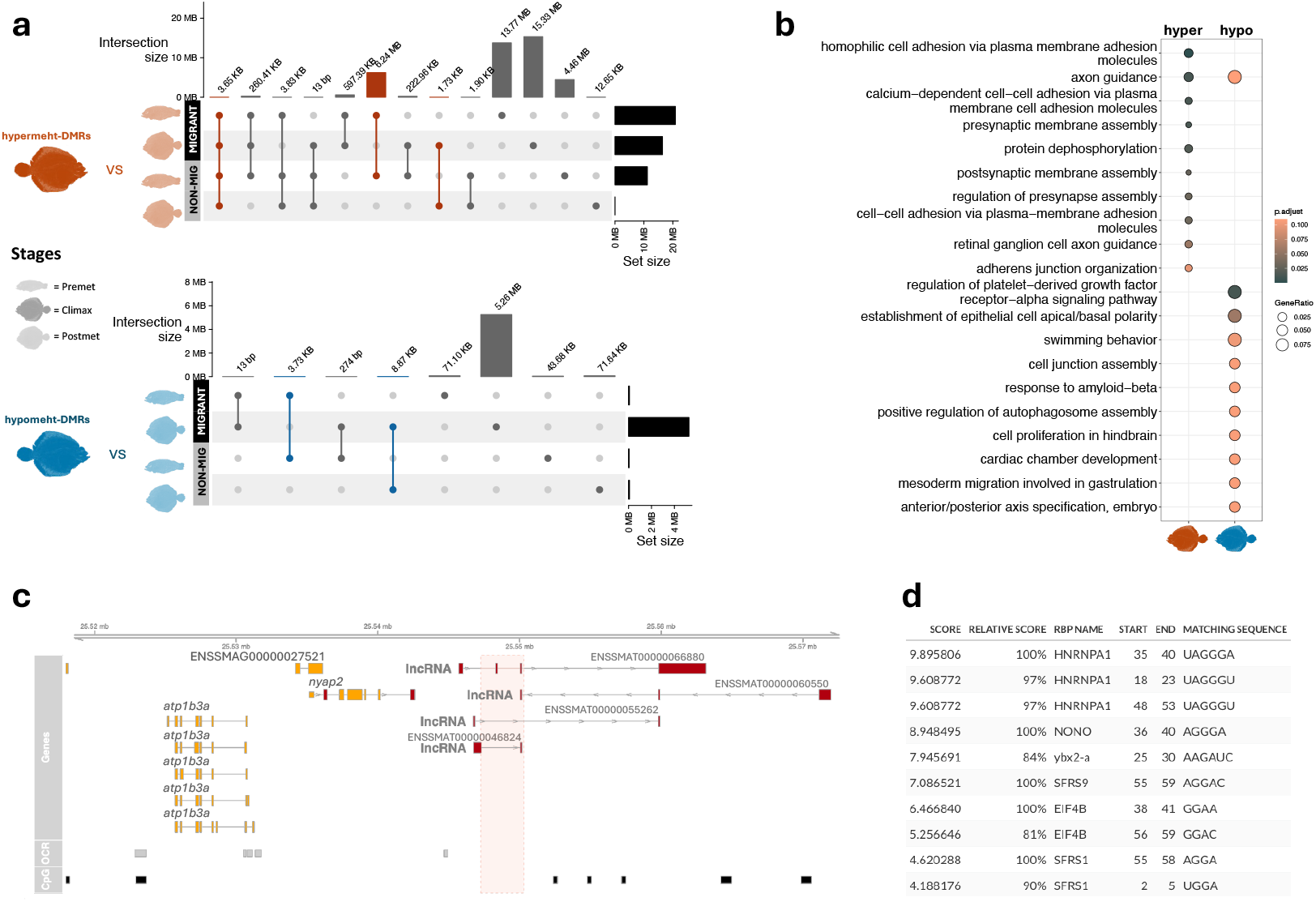
A conserved methylation pattern was observed in the migrating and non-migrating turbot eyes throughout the developmental stages. (a) UpSet plot showing regions of hypermethylation (orange) and hypomethylation (blue) at the culmination of metamorphosis, with a comparison between the climax and the pre- and post-metamorphosis stages for migrating (black panel) and non-migrating (grey panel) turbot eyes. The orange and blue lines highlight the regions that exhibit a similar methylation profile in both eyes. Icons representing the pre-metamorphic, climax, and post-metamorphic turbot stages, based on original photographs, are provided to illustrate the contrast between the stages. (b) Gene Ontology (GO) enrichment analysis of selected gene sets from the UpSet plot. Biological processes of genes that are in proximity to the hypermethylated (orange) or hypomethylated (blue) regions at the climax of metamorphosis, and with the same methylation pattern in both migrating and non-migrating eyes. (c) Visualization of a genomic region, which depicts a methylation peak at the climax of metamorphosis in both eyes. The red-shaded area indicates the location of DMR. The upper track illustrates the genomic localization, followed by gene annotation, which has been directly downloaded from ENSEMBL Biomart. The lowermost tracks illustrate open chromatin regions (OCR) from the NCBI GEO repository, with the series accession number of GSE215396, and CpG islands (CGI) annotated according to the Garden-Gardiner and Frommer algorithm. (d) Protein interactome predicted by linc2function software from the sequence of lncRNAs overlapping with the region of interest.

**Figure 2.**
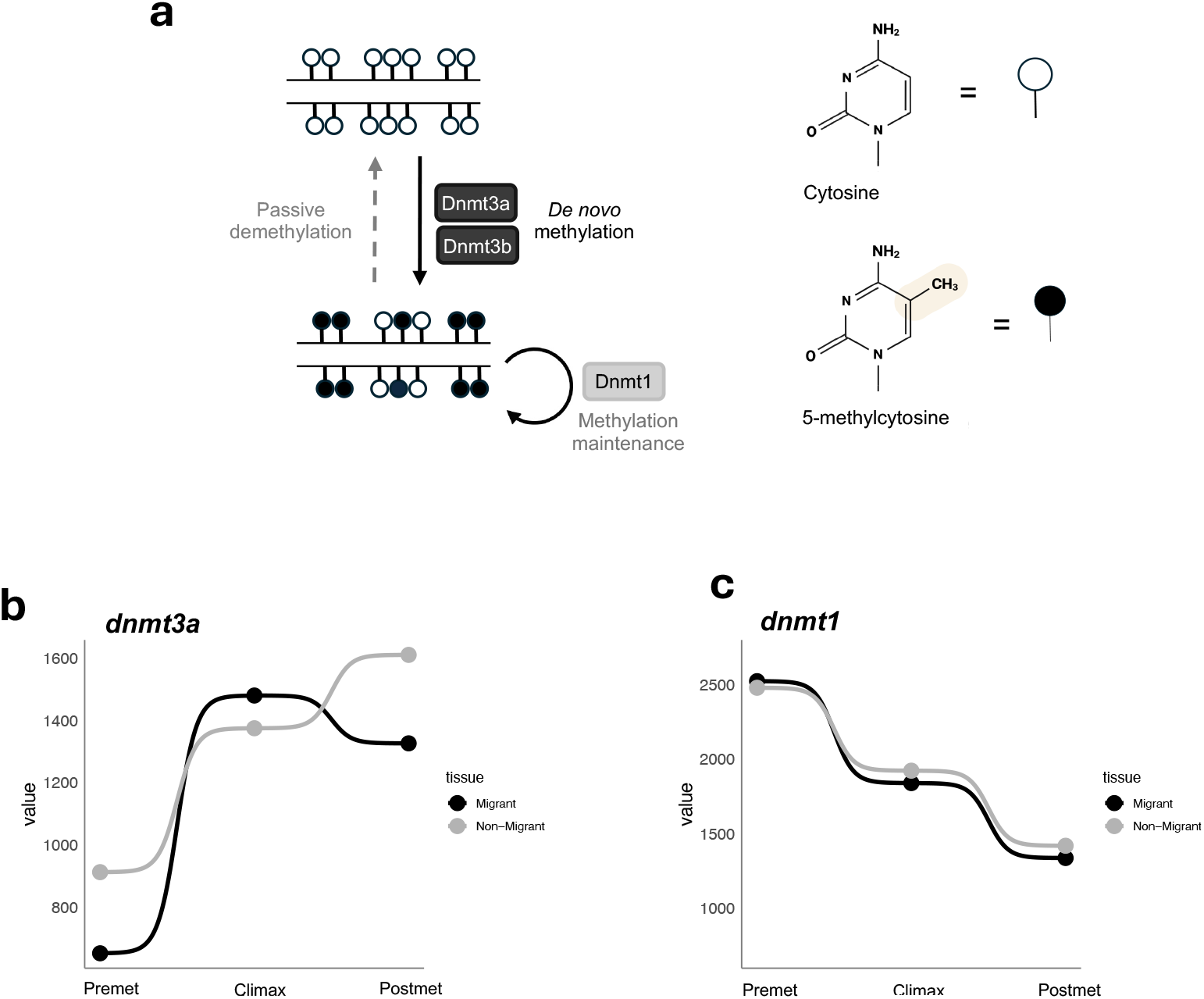
Dnmt1 and Dnmt3 in DNA methylation and gene expression across developmental stages in turbot. (a) Schematic representation of the functions of Dnmt1 and Dnmt3 in maintaining and establishing DNA methylation patterns. Unmethylated and methylated CpG dinucleotides are represented by empty and filled circles, respectively. Created with BioRender (https://biorender.com). (b) Normalized average counts of *dnmt3a* gene replicates in the three developmental stages of the migrating (black) and non-migrating (grey) eye. (c) Normalized average counts of *dnmt1* gene replicates at the three developmental stages of the migrating (black) and non-migrating (grey) eye.

One of the most notable visual adaptations observed in the flatfish as they transition to a benthic lifestyle, is their ability to see in low-light conditions. The expression of the principal isoforms expressed in photoreceptor cells, cones and rods, involved in the phototransduction cascade ***Figure 3***a, was examined focusing on the expression of a panel of cell-type marker genes (***Lamb, 2013***). Results showed a noticeable pattern wherein the isoforms expressed in cones exhibited a notable decline in expression during the postmetamorphic stage ***Figure 3***b. Conversely, the expression profile of the isoforms expressed in rods, which are responsible for scotopic vision, increased throughout the stages ***Figure 3***c. However, no significant correlation was identified between methylation and gene expression in the phototransduction cascade genes, neither in the promoter region nor in the first exons and introns. Furthermore, an anomalous ratio was observed in the number of CpG islands present among the set of genes implicated in the phototransduction cascade (Chi-squared = 19.5; p-value = 9.75e-06). This suggests that the regulation of this set of genes may not be dependent on methylation (***Tian et al., 2022***). Conversely, differential methylation was observed in the second exon of the *edem1* gene, which is crucial for the correct folding of rod opsin, across developmental stages but not between eyes ***Figure 3***d. Furthermore, a correlation was observed between gene expression and stages of development ***Figure 3***e.

**Figure 3.**
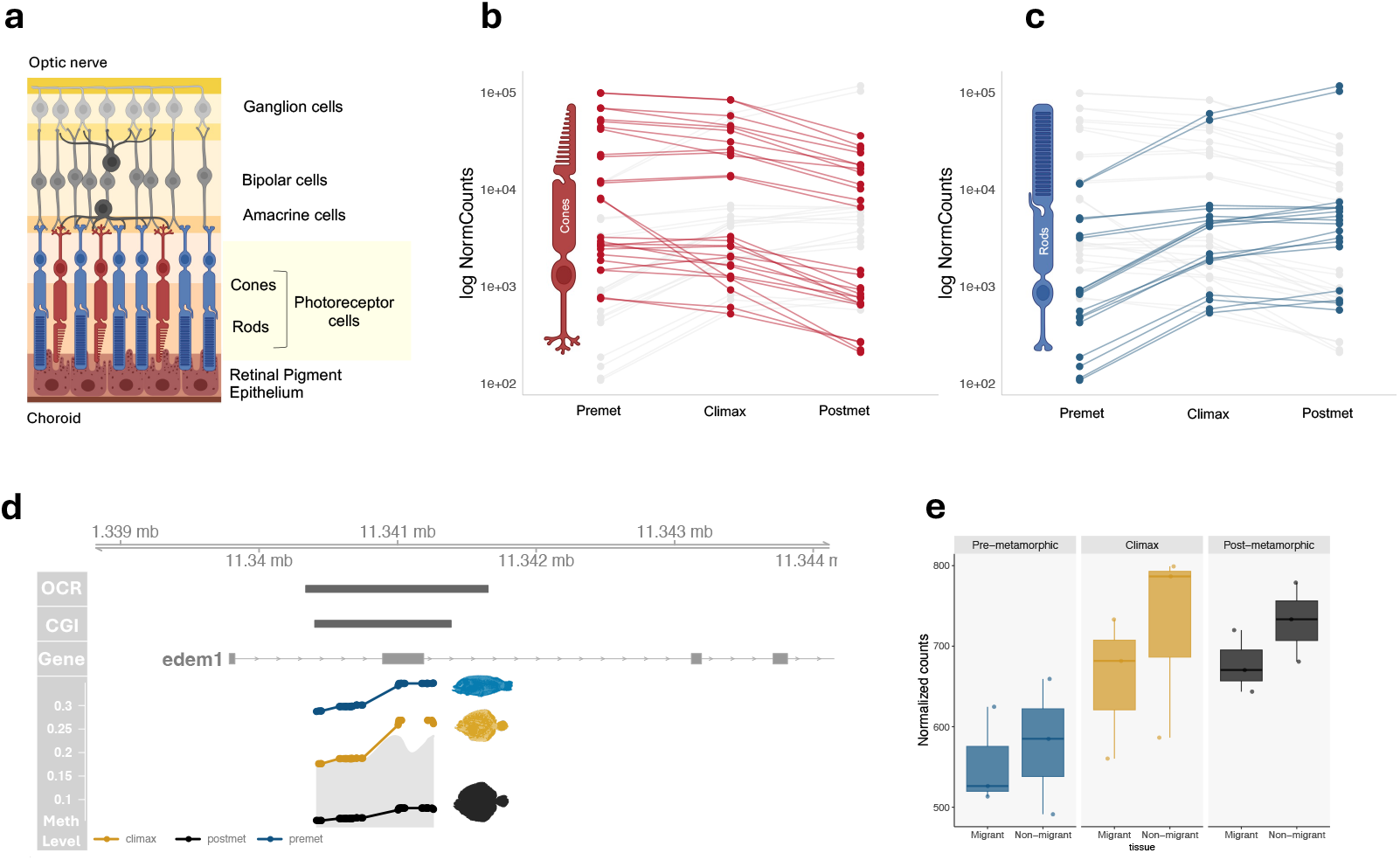
Epigenetic and gene expression profiles of genes associated with the phototransduction cascade. (a) Schematic representation of the retina, with an indication of the location of the photoreceptor cells, cones and rods. Created with BioRender (https://biorender.com). (b) Expression profiles (log of normalized counts) of genes involved in the phototransduction cascade throughout development in the migrating and non-migrating eyes, with cone-specific genes highlighted in red (pde6c, opn1sw1, gnb3a, gnb3b, cnrg, opn1sw2, opn1lw1, rh2-a1, rh2-b1 and rh2-b2) (***Lamb, 2013***). (c) Expression profiles (log of normalized counts) of genes involved in the phototransduction cascade throughout development in the migrating and non-migrating eyes, with rods-specific genes highlighted in blue (pde6a, cnga1a, gnb1, rh1, pde6g, gngt1, slc24a1, gnb1b, gnat1 and pdpr). (d) Visualization of the edem1 gene in the genomic context. The OCR track represents open chromatin regions, while the CGI track depicts the CpG islands that have been annotated using the Garden-Gardiner and Frommer algorithm. Colored tracks illustrate methylation levels at the pre-metamorphic (blue), climax of metamorphosis (yellow) and post-metamorphic (black) stages. (e) Expression profile (normalized counts) of the edem1 gene throughout turbot development in the migrating and non-migrating eye. Premet, climax and postmet labels correspond to pre-metamorphic, climax of metamorphosis and post-metamorphic stages, respectively.

### DNA methylation-mediated ocular migration across metamorphosis

The migration of one eye to the opposite side of the body is one of the most distinctive changes during flatfish metamorphosis. To investigate genome-wide DNA methylation differences in turbot we examined differential methylated regions (DMRs) between the migrating and non-migrating eye within the same developmental stage. A greater number of DMRs between the two eyes at the post-metamorphic stage were identified, with 51, 2921, and 11,260 DMRs in the pre-metamorphic, climax, and post-metamorphic stages, respectively ***Figure 4***a. Moreover, no significant differences were observed in genomic distribution of these DMRs for each stage, with a notable high proportion of them located in promoter and exonic regions in the three stages ***Figure 4***b. Gene ontology (GO) term enrichment analyses for the genes associated with the DMRs identified terms mainly related to neural fate, and compatible with ocular development, with the most notable being the negative regulation of canonical Wnt signaling pathway ***Figure 4***c as it plays an important role in the retinal vascular development (***Fujimura, 2016***).

**Figure 4.**
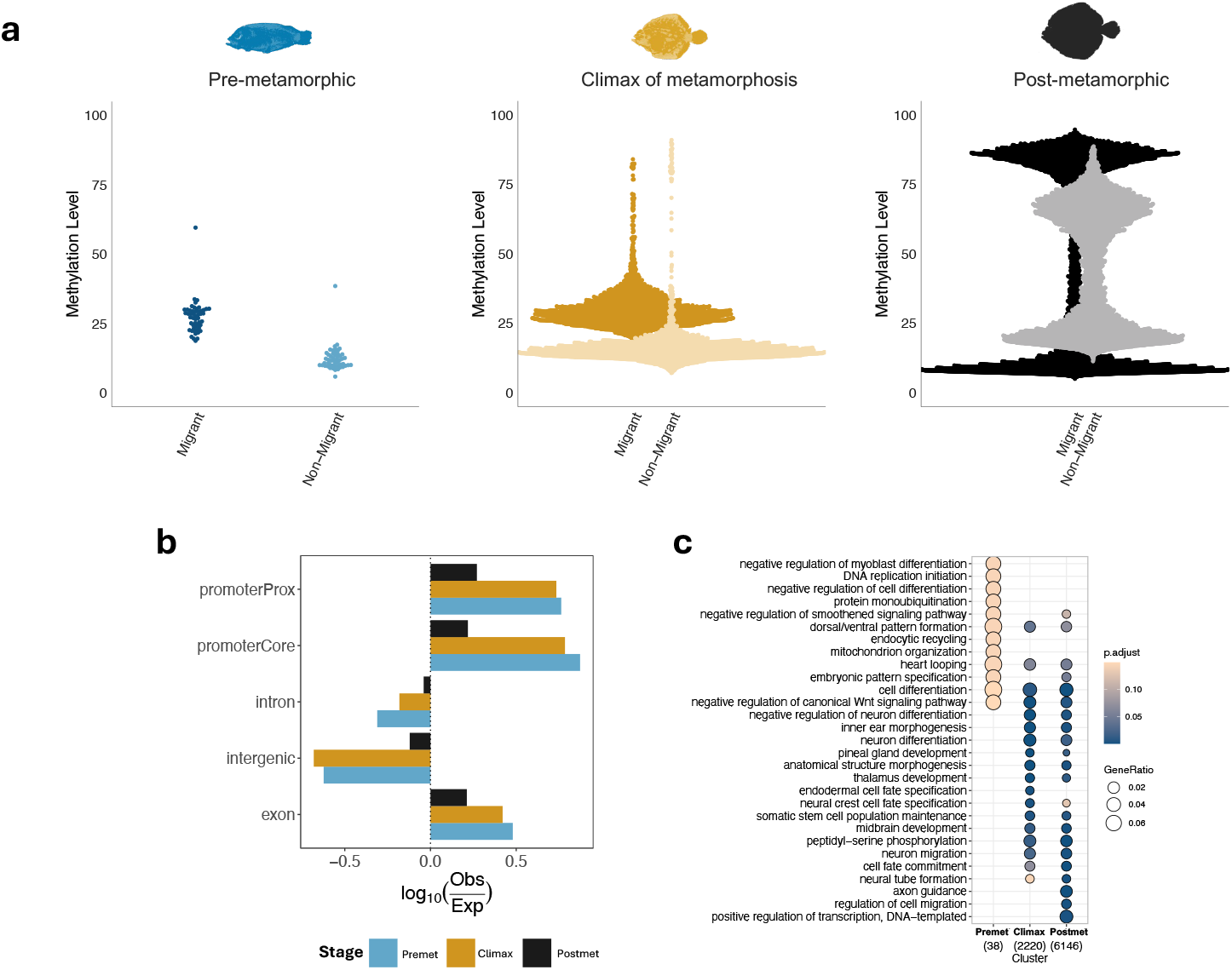
Comparative analysis of migrating and non-migrating eye methylation levels in the pre-metamorphic, climax and post-metamorphic stages of turbot. (a) Beeswarm plot illustrating methylation levels of all DMRs identified in migrating and non-migrating eye at the pre-metamorphic (blue), climax of metamorphosis (yellow) and post-metamorphic stages (black). Each data point represents a DMR identified through contrast. (b) Enrichment plot across genome annotation of the DMRs identified by comparison of the migrating and non-migrating eye at the three key stages of turbot development. (c) Gene ontology (GOs) enrichment analysis of the DMRs identified between the two eyes at the three developmental stages in turbot. The numbers on the X-axis represent the total number of genes in each stage.

In order to investigate the relationship between DNA methylation and gene expression during the eye migration process, we compared methylation and expression changes obtained from the comparison between eyes at different developmental stages ***Figure 5***a. At the pre-metamorphic stage, all identified DMRs were hypermethylated in the migrating eye, with a positive correlation evident in approximately half of the cases. A hypermethylation trend in the migrating eye persisted during the climax stage, accompanied by a notable increase in the number of identified DMRs between the two eyes. Moreover, 1,237 of these DMRs exhibited a negative correlation with gene expression, while 1,399 DMRs demonstrated a positive correlation. Conversely, the number of hypermethylated DMRs began to compensate in the migrating (4,986) and non-migrating (4,753) eye during the post-metamorphic stage. The genomic features with overlapping DMRs that are negatively and positively correlated with gene expression were also examined. No significant differences were identified in the genomic distribution of DMRs ***Figure 5***b.

**Figure 5.**
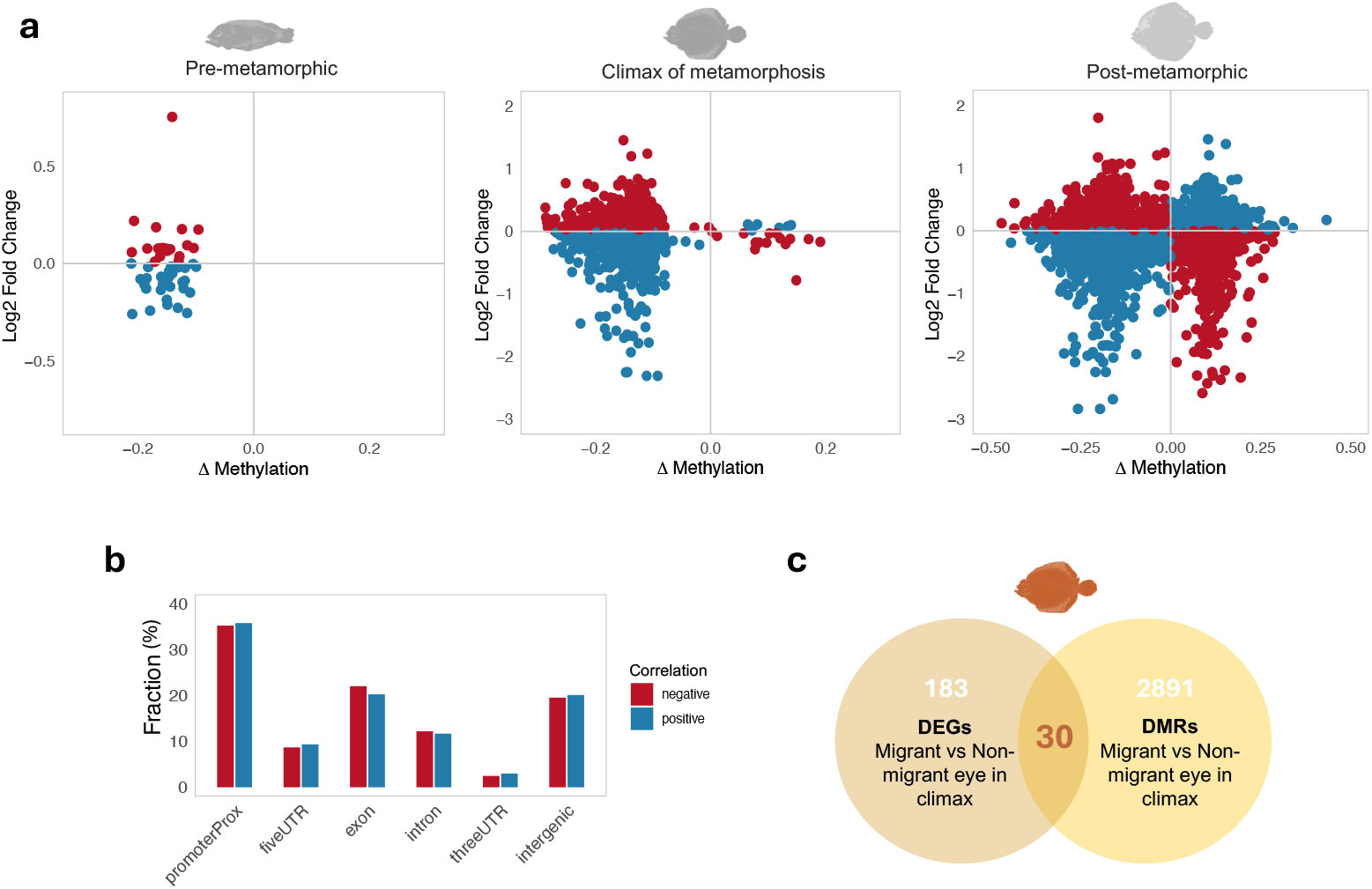
Correlation between methylation and gene expression of genes involved in flatfish eye migration. (a) Scatter plot representing the log2 fold change versus methylation increase identified between the migrating and non-migrating eye at the pre-metamorphic, climax of metamorphosis and post-metamorphic stages. The red dots indicate negatively correlated DMRs with respect to gene expression, whereas the blue dots represent positive correlated DMRs. (b) Bar chart illustrating the proportion of all identified DMRs that are positively (blue) or negatively (red) correlated with annotated genomic regions. (c) Venn diagram illustrating the overlap between differentially expressed genes (DEGs) and differentially methylated regions (DMRs) between migrating and non-migrating eyes at the climax stage of metamorphosis.

In terms of gene expression, we identified 213 differentially expressed genes (DEGs) between the migrating and non-migrating eyes during climax stage. 30 of these colocalized with differentially methylated regions (DMRs) between the migrating and non-migrating eye during this stage ***Figure 5***c. Among the 30 differentially expressed and methylated genes between eyes in the climax, two transcription factors were of particular interest due to their functions in the development and maintenance of retinal ganglion cells. The transcription factor *eomesa* exhibited a region in the final exon with a 11% difference in methylation between the migrating and the non-migrating eyes. Furthermore, this region maps to a CpG island and a region of accessible chromatin ***Figure 6***a. Conversely, the transcription factor *tbr1b* exhibited a methylation discrepancy of approximately 14% between the migrating and non-migrating eyes, located within the first exon and aligning with a CpG island ***Figure 6***a. An examination of the expression profiles of these genes revealed maximum expression at the climax in the migrating eye while in the non-migrating eye there was a decline in expression after the pre-metamorphic stage ***Figure 6***b. As previously described by ***Kölsch et al***. (***2021***), the *eomesa+* RGCs are primarily situated in the ventral region of the retina and project multiple arborization fields, extending to the tectum in zebrafish. However, the distribution of the *tbr1b+* RGCs is limited to the retina, with direct innervation to the intermediate tectal layers ***Figure 6***c.

**Figure 6.**
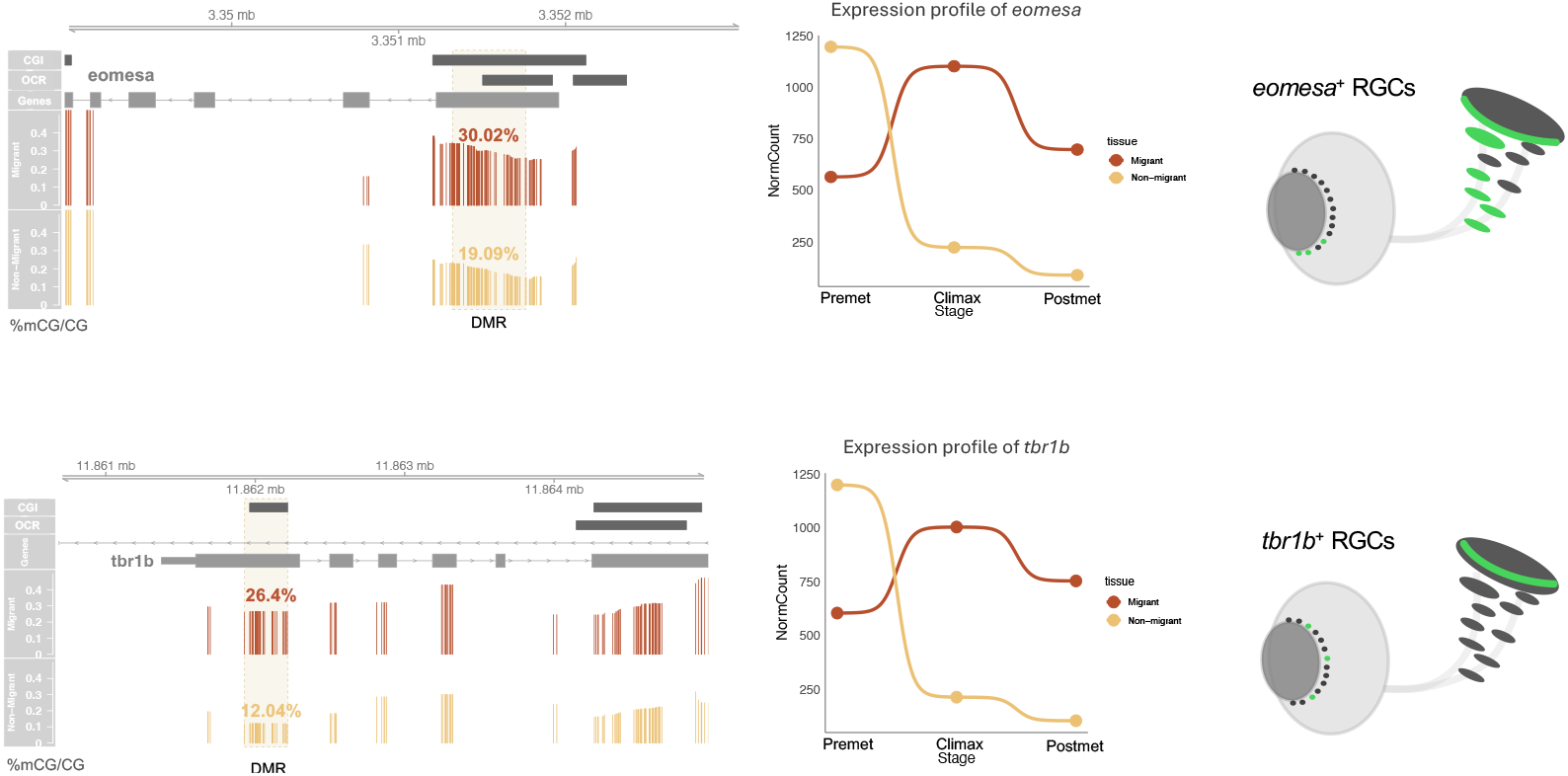
Methylation and expression profiles of the transcription factors involved in the development of retinal ganglion cells (RGCs). (a) Genomic view of the *eomesa* and *tbr1b* gene. The two first tracks at the top illustrate annotated CpG islands (CGI) and open chromatin regions (OCR). The two tracks at the bottom illustrate the methylation levels per CpG dinucleotide at the climax stage of metamorphosis in the migrating (orange, top) and non-migrating (yellow, bottom) eyes. The differentially methylated region between the two eyes is indicated by shading. (b) Gene expression profile (mean normalized counts) of the *eomesa* and *tbr1b* genes in the migrating (orange) and non-migrating (yellow) eye throughout turbot development. (c) Schematic representation of the soma distribution in the retina and the projection pattern of *eomesa+* RGC (black) and *tbr1b+* RCG (green) in zebrafish. The image has been modified from that used in ***Kölsch et al***. (***2021***). The TF+ RGCs for each gene are shown in green. The *eomesa+* RGCs are distributed in the ventral part of the retina and innervate multiple arborization fields, extending down to the tectum. The distribution of the *tbr1b+* RGCs is localized in the retina and directly innervate the intermediate tectal layers.

## Discussion

The migration of one eye to the opposite side of the body represents a distinctive adaptation that enables flatfish to adopt a juvenile benthic lifestyle, thereby optimizing their vision while lying on the ocean floor. The understanding of the differences in gene regulation between the migrating and the non-migrating eye is essential to shed light on the morphological remodeling that flatfish undergo during metamorphosis. DNA methylation plays a pivotal role in regulating gene expression, which is vital for the successful completion of metamorphosis. In this study, we investigated the methylation profiles of the migrating and non-migrating eye during three main stages of turbot development. We analyzed the correlation between methylation pattern and gene expression, identifying differentially methylated regions (DMRs) specific to each eye for each stage. This is the first study that examines the specific DNA methylation profile in the eyes of a flatfish species. Although it would be expected that DNA demethylation would occur during metamorphosis in response to a potential reprogramming and recapitulation to earlier developmental stages, a significantly high number of DMRs were found to be hypermethylated at the climax stage in both eyes when compared with any other stage. These findings are consistent with those of studies conducted on amphibian metamorphosis (***Kyono et al., 2020***). It has been demonstrated that thyroid hormone can induce transcription of the *dnmt3a* gene in *Xenopus* (***Kyono et al., 2016***), which is responsible for establishing the global methylation pattern through de novo methylation (***Okano et al., 1999***). As in amphibians (***Tata, 1998***), metamorphosis in flatfish is completely T3-dependent (***de Jesus et al., 1993***). In this study, we also demonstrated that *dnmt3a* gene expression is increased during the climax of metamorphosis in turbot. Consequently, stagespecific hypermethylated DMRs were identified at the climax stage in comparison to other developmental stages. By examining conserved DNA methylation patterns in both eyes in response to benthic adaptation, we identified three regions that exhibited a methylation peak at the climax of metamorphosis in migrating and non-migrating eyes. One of these regions exhibited colocalization with lncRNA sequences. Long noncoding RNAs (lncRNAs) play a key role in genome regulation, exerting their influence through both cis and trans mechanisms (***McKiernan and Greene, 2015; Kulczyńska and Siatecka, 2016***). The neighboring proteincoding genes to these lncRNAs of interest may have potential local action (***Vance and Ponting, 2014; Engreitz et al., 2016***). Interestingly, the genes located in closest proximity to their respective synthesis sites are the *atp1b3b* gene, which serves to prevent the dysregulation of ion channel activity, thus averting the potential disruption of the ionic balance within retinal ganglion cells; and the *nyap2a* gene, which plays a role in the morphogenesis of neuronal projections. Furthermore, the identification of specific protein binding motifs within these particular lncRNAs revealed their capacity to interact and form complexes with the RNA binding proteins (RBPs) ELAVL1, NONO, and HNRPA1B, previously shown to be involved in eye development (***Dash et al., 2016***), specifically in lens formation (***Coomson et al., 2022***), neural retina development (***Neant et al., 2011***), and regulation of neuronal-specific splicing (***Jia et al., 2019; Gillentine et al., 2021***). Although it has been established that the flatfish eye does not undergo developmental changes in the ocular layers or lens during metamorphosis (***Villegas et al., 1997***), other studies have reported an overexpression of genes involved in lens development during this stage (***Guerrero-Peña et al., 2024***). Therefore, our findings suggest that the adaptation to benthic environment may also be regulated by lncRNAs, possibly through interactions with RBPs that influence lens-related gene expression, even in the absence of physical changes in the lens structure. The development of a camouflage and ambush strategy is part of the flatfish transition from a pelagic existence during their larval stage to a benthic lifestyle during their juvenile phase (***Friedman, 2008; Schreiber, 2006***). This radical transition compels them to adapt morphologically and behaviorally in a remarkably brief period to entirely novel environmental conditions. With regards to vision, the photic environment of the water undergoes rapid changes with depth (***Davies et al., 2012; Hauser and Chang, 2017***). Consequently, flatfish develop effective scotopic vision during their juvenile phase due to to photoreceptor reorganization and alterations in opsin expression (***Wang et al., 2021; Sara Frau, Iñigo Novales Flamarique, Patrick W***. ***Keeley, Benjamin E***. ***Reese, 2017; Mader and Cameron, 2004***). The expression profile of cone and rod-specific opsin genes demonstrates a decrease in the former and an increase in the latter through the three developmental stages, which is consistent with what is expected based on flatfish settlement. However, no changes in DNA methylation were observed in the promoter region between stages. It is noteworthy that the majority of genes involved in the phototransduction cascade do not possess a CpG island in their promoter region. While previous studies have underscored the function of DNA methylation in the control of photoreceptor gene expression (***Merbs et al., 2012***), the epigenetic modulation of cone and rod opsins remains unclear (***Rao et al., 2011; Lu et al., 2023***). The absence of CpG islands and the limited alterations in DNA methylation observed in the promoters of phototransduction cascade genes suggest that their regulation may not be dependent on DNA methylation in turbot. It has been estimated that more than half of the genes present in the turbot genome colocalize with a CpG island in its promoter region. Thus, it is well-known that CpG islands play a role in transcriptional regulation, particularly in the case of constitutively expressed and tissue-specific genes. However, it has been demonstrated that orphan CpG islands, which do not colocalize with promoters, may function as unannotated distal promoters (***Maunakea et al., 2010; Illingworth et al., 2010***). It is therefore plausible that the cone and rod-specific genes responsible for visual adaptation to low-light conditions have mostly unidentified distal promoters, being possible a DNA methylation-dependent regulation. Our results support the hypothesis that cones and rods undergo a change during flatfish metamorphosis, as evidenced by alterations in the expression and DNA methylation of edem1, a gene essential for rod opsin biosynthesis (***Kosmaoglou et al., 2009; Athanasiou et al., 2014***). Furthermore, edem1 appears to be subjected to regulation by DNA methylation mechanisms. In addition to the modifications in the visual system, flatfish exhibit a migration of one eye to the opposite side of the body. The aim of this study was also to investigate potential differences in DNA methylation between the two eyes (migrating and non-migrating eyes). The transcription factors *eomesa* and *tbr1b* are of interest due to their hypermethylation and overexpression profiles at the climax of metamorphosis in the migrating eye. The *eomesa* gene has been demonstrated to regulate the process of retinal ganglion cell (RGC) differentiation and optic nerve development (***Mao et al., 2008***), while *tbr1b* has been characterized as a prerequisite for the laminar specification of RGC dendrites (***Liu et al., 2018***). The primary function of retinal ganglion cells is to transmit visual information from the retina to the brain through the optic nerve. The axonal projections from these cells terminate in the arborization fields innervating the tectum (***Robles et al., 2014***). In zebrafish, *tbr1b+* RGCs have been identified in the retina with axons that directly innervate the intermediate tectal layer. In contrast, *eomesa+* RGCs in the retina have been observed to innervate multiple extratectal areas (***Kölsch et al., 2021***). The function of these genes in neuronal differentiation may be crucial for the remodeling of the optic nerve during metamorphosis, as well as for the proper formation and projection of axons to the brain, which is essential for visual function. Although the vestibulo-ocular reflex has been the subject of study in flatfish (***Graf and Baker, 1983; Graf et al., 2001; Graf and Baker, 1990; Graf et al., 1997***), the changes in axonal projections that occur during metamorphosis have not been fully described. However, previous studies have noted a shortening of the optic nerve in the migrating eye9 and a reduction in the size of the migrating-eye optic tectum at the climax of metamorphosis (***Briñón et al., 1993***). Flatfish undergo T3-driven metamorphic remodeling, which necessitates the adaptation of their vision to the restricted light conditions that prevail on the seafloor. The T3 hormone, which induces *de novo* methylation via *dmnt3a*, promotes differential hypermethylation during the climax of metamorphosis. In contrast, the observed expression of genes involved in the phototransduction cascade indicates a remodeling of the opsin pool, which appears to be adapted to low-light conditions. Nevertheless, it remains unclear whether the regulation of these genes is methylation-dependent. In the specific comparative study between migrating and non-migrating eyes during metamorphosis, we describe the activation of pathways involved in ocular development and retinal regeneration. Our results demonstrated distinctive expression and methylation patterns of the transcription factors *eomesa* and *tbr1b*, suggesting a significant reorganization of visual projections between the migrating and non-migrating eye. These changes suggest that RGCs engaged in substantial activity during metamorphosis in the migrating eye, likely associated with the transmission of the visual signal to the brain.

This study provides novel insights into the epigenetic mechanisms underlying flatfish metamorphosis, particularly regarding eye migration and visual adaptation. For the first time, we have characterized the DNA methylation profiles specific to migrating and non-migrating eyes during turbot development, revealing stage-specific hypermethylation during metamorphosis climax that is regulated by T3-induced *dnmt3a* expression. Our findings highlight the role of lncRNAs in benthic adaptation through potential interactions with eye development-related RBPs, despite the absence of structural changes in the lens. Furthermore, we identified differential methylation patterns in key transcription factors (*eomesa* and *tbr1b*) between migrating and non-migrating eyes, suggesting their crucial role in optic nerve remodeling and retinal ganglion cell reorganization during metamorphosis. These discoveries advance our understanding of the complex molecular mechanisms governing flatfish metamorphosis and provide a starting point for future studies investigating the relationship between epigenetic regulation and morphological adaptation in marine species.

## Methods and Materials

### Sampling of turbot tissues and stages

Newborn turbots (*Scophthalmus maximus*) were obtained from Insuiña SL, a fish farm part of the Nueva Pescanova Group (Pontevedra, Spain). Fish were reared in accordance with standard commercial production conditions at 18°C. The offspring of a single breeding pair was sampled at three different developmental stages: pre-metamorphic stage (13 days post-fertilization, dpf), metamorphic climax (21 dpf) and post-metamorphic stage (44 dpf), with n = 3 independent biological replicates per stage. The metamorphic stages were defined in accordance with the morphological criteria established by Al-Maghazachi and Gibson (***Al-Maghazachi and Gibson, 1984***) and Suarez-Bregua (***Suarez-Bregua et al., 2020***). The animals were euthanized using a lethal dose of MS-222 (Sigma-Aldrich, USA) (500 mg/L for 30 min). Subsequently, the eyes of each individual were extracted with the aid of a Leica M165FC stereomicroscope and stored at – 80 °C. The number of fish used in all experiments was calculated to ensure that the results were statistically significant and reliable, while adhering to the minimum sample size required for statistical power. All procedures were conducted in accordance with the ethical standards set forth by the Institutional Committee for the Care and Use of Animals of the IIM-CSIC Institute (ES360570202002/17/FUN.1/BIOLAN.08/JRM), in compliance with the guidelines established by the National Advisory Committee for Research with Laboratory Animals and authorized by the Spanish Authority (RD53/2013). The research was conducted in accordance with the European directive (2010/63/EU) for the protection of animals used in scientific experiments.

### DNA isolation, libraries preparation and sequencing

The eyes of each specimen were extracted for each key developmental stage, identifying the eye that underwent migration and the one that did not. Three independent biological replicates of each condition were subjected to mechanical homogenization and treated with DNAse-free RNAse (final concentration 2 µg/µl) and protease. DNA was extracted and purified using the QIAGEN Genomic DNA kit (Qiagen, Germany), in accordance with the manufacturer’s instructions. Subsequently, Reduced Representation Bisulfite Sequencing (RRBS) libraries were prepared in accordance with the quantification of the 18 extracted DNA samples using a Qubit fluorometric assay (ThermoFisher Scientific, USA). The Diagenode Premium RRBS kit (Diagenode Inc., USA) was used in accordance with the manufacturer’s instructions. The DNA from all samples was subjected to bisulfite conversion by MspI enzyme digestion, resulting in a total of 100 ng of DNA targeted for CCGG sites. The resulting DNA fragments were purified using AMPure XP Beads (Beckman-Coulter, USA), and their concentration was determined using a Qubit High Sensitivity assay (ThermoFisher Scientific, USA). Thereafter, the libraries were subjected to sequencing on an Illumina HiSeq2000 platform, utilizing 50 bp single-end reads.

### Determination of eye and stage-specific differentially methylated regions

The quality of the sequenced data was evaluated using the FastQC v0.12.0 software, and adapters were removed using the TrimGalore v0.6.10-0 tool (***Krueger et al., 2023***). The efficacy of the CpG bisulfite conversion process was validated through the examination of spike-in controls. The RRBS sequenced reads were subsequently aligned to the turbot reference genome (ASM1334776v1) using BSMAP v2.90 (***Xi and Li, 2009***), with methylation analysis conducted using the methratio.py script, which is included in the BSMAP software. A differential methylation analysis was conducted across the three stages using the DSS v2.50.1 R package (***Wu et al., 2015***). Two types of pairwise comparisons were conducted using the Wald test with the DMLtest function with the smoothing parameter set to TRUE, to determine methylation difference at each locus. Then, these results were loaded to callDMR function in order to determine the length of the regions. Differentially methylated regions (DMRs) were determined by a Wald test with a p-value threshold of 0.05 and a minimum CG content of 10. Two pairwise comparisons were conducted: one based on turbot developmental stages (pre-metamorphosis, climax of metamorphosis and post-metamorphosis) and other based on the type of eye tissue (migrating and non-migrating eye) within each developmental stage. In order to study concordant methylated regions that exhibit similar methylation patterns in both migrating and non-migrating eyes throughout the various developmental stages, the hypermethylated and hypomethylated regions matching those observed in both eyes during the climax stage were studied. Three distinct datasets were extracted using the UpSet function from the ComplexHeatmap v2.18.0 R package (***Gu et al., 2016; Gu, 2022***): (1) a set of DMRs at the climax stage vs. the pre-metamorphic stage in both eyes; (2) a set of DMRs at the climax stage vs. the post-metamorphic stage in both eyes; (3) a set of DMRs at the climax stage in both eyes. The tissue-specific methylation levels were visualized in a beeswarm plot.

### RNA isolation and sequencing

Total RNA was extracted and purified from all collected samples using the RNeasy Mini Kit (Qiagen, Germany), in accordance with the manufacturer’s instructions. Following verification of the integrity of the extracted RNA using an Agilent 2100 Bioanalyzer (Agilent Technologies, USA), samples with an RNA integrity number (RIN) of 8 or higher were selected for library preparation. The DNBSEQ stranded mRNA libraries were sequenced on a DNBseq™ platform (BGI Genomics, Hongkong), resulting in 100 base pair (bp) paired- end reads for each sample.

### Identification of differential gene expression across stages and tissues

The quality of all reads was evaluated using the FastQc v0.12.0 tool. Only high-quality raw reads were aligned against the most recent turbot genome (ASM318616v1) using the STAR v2.7.0e (***Dobin et al., 2013***) alignment software. The HTSeq software (version 0.10.0) (***Anders et al., 2015***) was used to assign mapped reads to a single annotated gene. The count matrices were normalized using median-of-ratio methods with the DESeq2 v1.42.1 R package (***Love et al., 2014***). Pairwise comparisons of differential gene expression were conducted at both the stage and tissue levels with DESeq2 v1.42.1 R package. Genes that met the criteria of a log2 fold change greater than 1 in absolute values and a p-value less than 0.05 were identified as differentially expressed (DEG). The expression profiles of genes of interest throughout development were examined using bumplots and boxplots generated in R with ggbump v0.1.0 and ggplot2 v.3.5.1 packages, with all cases representing the normalized counts.

### Correlation between DNA methylation and gene expression

In order to examine the relationship between DNA methylation, gene expression and gene ontology (GO enrichment analysis), we first associated the genomic coordinates of the regions identified as differentially methylated with the annotated genes or the closest gene present within the turbot genome. Consequently, the genomation v1.34.0 R package (***Akalin et al., 2015***) was used for the annotation of gene parts. The correlation between DNA methylation and gene expression was evaluated by plotting the mean methylation level of the gene region against the corresponding log2 fold change values in a scatter plot. A GO enrichment analysis was conducted by computing functional profiles for each gene cluster using the compareCluster function from the ClusterProfiler (***Yu et al., 2012***) R package v3.14.3. To create the custom background, a specialized functional annotation of the turbot genome created with Sma3s (***Muñoz-Mérida et al., 2014; Casimiro-Soriguer et al., 2017***) v2.1 was used.

### Genomic feature annotation

We used the GenomicFeatures (***Lawrence et al., 2013***) v1.54.4 R package to create a gene model from the turbot genome annotation, dividing the genome into promoters, exons, introns, and intergenic regions. Each identified DMR was assigned to one of these features, and the distributions was calculated by overlap. We calculated the DMR enrichment for each genomic feature and performed Chi-squared tests to compare the observed and expected overlap of DMRs in each comparison group using the calcExpectedPartitions function from the GenomicDistributions (***Kupkova et al., 2022***) v1.10.0 R package. CpG islands were predicted using the Gardiner-Garden and Frommer algorithm (***Gardiner-Garden and Frommer, 1987***), based on the following criteria: (1) GC content of 50% or more, (2) length greater than 200 base pairs, and (3) an observed/expected ratio greater than 0.6.

### Genomic browser

We visualized the genomic data for the genes of interest using the Gviz (***Hahne and Ivanek, 2016***) v1.46.1 R package. Genome browser tracks were created by incorporating genomic annotation features, chromatin peak data from the NCBI Gene Expression Omnibus repository (GSE215396 ***Guerrero-Peña et al***. (***2023***) series accession number).

## Acknowledgments

Additional information can be given in the template, such as to not include funder information in the acknowledgments section. This work was made using a template from *eLife* journal provided under the CC BY 4.0 license with some minor style changes.

## References

Akalin A, Franke V, Vlahoviček K, Mason CE, Schübeler D. Genomation: a toolkit to summarize, annotate and visualize genomic intervals. Bioinformatics (Oxford, England). 2015 apr; 31(7):1127–1129. https://pubmed.ncbi.nlm.nih.gov/25417204/, doi: 10.1093/BIOINFORMATICS/BTU775.

Al-Maghazachi SJ, Gibson R. The developmental stages of larval turbot, Scophthalmus maximus (L.). Journal of Experi-mental Marine Biology and Ecology. 1984; 82:35–51.

Anders S, Pyl PT, Huber W. HTSeq-A Python framework to work with high-throughput sequencing data. Bioinformatics. 2015; doi: 10.1093/bioinformatics/btu638.

Athanasiou D, Bevilacqua D, Aguila M, McCulley C, Kanuga N, Iwawaki T, Paul Chapple J, Cheetham ME. The co-chaperone and reductase ERdj5 facilitates rod opsin biogenesis and quality control. Human molecular genetics. 2014; 23(24):6594– 6606. doi: 10.1093/hmg/ddu385.

Bogdanović O, Long SW, Heeringen SJV, Brinkman AB, Gómez-Skarmeta JL, Stunnenberg HG, Jones PL, Veenstra GJC. Temporal uncoupling of the DNA methylome and transcriptional repression during embryogenesis. Genome Research. 2011; 21:1313–1327. doi: 10.1101/gr.114843.110.21.

Bogdanović O, Heeringen SJV, Veenstra GJC. The epigenome in early vertebrate development. Genesis. 2017; 50(March):192–206. doi: 10.1002/dvg.20831.

Bolstad K, Novales Flamarique I. Photoreceptor distributions, visual pigments and the opsin repertoire of Atlantic halibut (Hippoglossus hippoglossus). Scientific Reports. 2022; 12(1):1–24. https://doi.org/10.1038/s41598-022-11998-9, doi: 10.1038/s41598-022-11998-9.

Briñón JG, Médina M, Arévalo R, Alonso JR, Lara JM, Aijón J. Volumetric Analysis of the Telencephalon and Tectum During Metamorphosis in a Flatfish, the Turbot Scophthalmus maximus. Brain, Behavior and Evolution. 1993; 41(1):1–5. doi: 10.1159/000113819.

Casimiro-Soriguer CS, Muñoz-Mérida A, Pérez-Pulido AJ. Sma3s: A universal tool for easy functional annotation of pro-teomes and transcriptomes. Proteomics. 2017; doi: 10.1002/pmic.201700071.

Cheng SY, Leonard JL, Davis PJ. Molecular Aspects of Thyroid Hormone Actions. Endocrine Reviews. 2010 apr; 31(2):139. /pmc/articles/PMC2852208//pmc/articles/PMC2852208/?report=abstract https://www.ncbi.nlm.nih.gov/pmc/articles/PMC2852208/, doi: 10.1210/ER.2009-0007.

Coomson S, Aryal S, Lachke SA. The role of the RNA-binding protein Elavl1 in lens development. Investigative Ophthal-mology Visual Science. 2022; 63(7):652. https://iovs.arvojournals.org/article.aspx?articleid=2779506.

Dash S, Siddam AD, Barnum CE, Janga SC, Lachke SA. RNA Binding Proteins in Eye Development and Dis-ease: Implication of Conserved RNA Granule Components. Wiley interdisciplinary reviews RNA. 2016 jul; 7(4):527. /pmc/articles/PMC4909581//pmc/articles/PMC4909581/?report=abstract https://www.ncbi.nlm.nih.gov/pmc/articles/PMC4909581/, doi: 10.1002/WRNA.1355.

Davies WIL, Collin SP, Hunt DM. Molecular ecology and adaptation of visual photopigments in craniates. Molecular Ecology. 2012 jul; 21(13):3121–3158. https://onlinelibrary.wiley.com/doi/full/10.1111/j.1365-294X.2012.05617.x https://onlinelibrary.wiley.com/doi/abs/10.1111/j.1365-294X.2012.05617.x https://onlinelibrary.wiley.com/doi/10.1111/j.1365-294X.2012.05617.x, doi: 10.1111/J.1365-294X.2012.05617.X.

Dobin A, Davis CA, Schlesinger F, Drenkow J, Zaleski C, Jha S, Batut P, Chaisson M, Gingeras TR. STAR: Ultrafast universal RNA-seq aligner. Bioinformatics. 2013; 29(1):15–21. doi: 10.1093/bioinformatics/bts635.

Engreitz JM, Haines JE, Perez EM, Munson G, Chen J, Kane M, McDonel PE, Guttman M, Lander ES. Local regulation of gene expression by lncRNA promoters, transcription and splicing. Nature. 2016; 539(7629):452–455. https://pubmed.ncbi.nlm.nih.gov/27783602/, doi: 10.1038/NATURE20149.

Evans BI, Fernald RD. Retinal transformation at metamorphosis in the winter flounder (Pseudopleuronectes amer-icanus). Visual Neuroscience. 1993 nov; 10(6):1055–1064. https://www.cambridge.org/core/product/identifier/S0952523800010166/type/journal_article, doi: 10.1017/S0952523800010166.

Friedman M. The evolutionary origin of flatfish asymmetry. Nature. 2008; 454(7201):209–212. doi: 10.1038/nature07108.

Fujimura N. WNT/β-catenin signaling in vertebrate eye development. Frontiers in Cell and Developmental Biology. 2016 nov; 4(NOV):221674. https://www.frontiersin.org, doi: 10.3389/FCELL.2016.00138/BIBTEX.

Gardiner-Garden M, Frommer M. CpG Islands in vertebrate genomes. Journal of Molecular Biology. 1987 jul; 196(2):261–282. doi: 10.1016/0022-2836(87)90689-9.

Gillentine MA, Wang T, Hoekzema K, Rosenfeld J, Liu P, Guo H, Kim CN, De Vries BBA, Vissers LELM, Nordenskjold M, Kvarnung M, Lindstrand A, Nordgren A, Gecz J, Iascone M, Cereda A, Scatigno A, Maitz S, Zanni G, Bertini E, et al. Rare deleterious mutations of HNRNP genes result in shared neurodevelopmental disorders. Genome Medicine. 2021 ec; 13(1):31. /pmc/articles/PMC8056596//pmc/articles/PMC8056596/?report=abstract https://www.ncbi.nlm.nih.gov/pmc/articles/PMC8056596/, doi: 10.1186/S13073-021-00870-6.

Gomes AS, Alves RN, Rønnestad I, Power DM. Orchestrating change: The thyroid hormones and GI-tract development in flatfish metamorphosis. General and Comparative Endocrinology. 2015; 220:2–12. http://dx.doi.org/10.1016/j.ygcen.2014.06.012, doi: 10.1016/j.ygcen.2014.06.012.

Graf W, Baker R. Adaptive changes of the vestibulo-ocular reflex in flatfish are achieved by reorganization of central nervous pathways. Science (New York, NY). 1983; 221(4612):777–779. https://pubmed.ncbi.nlm.nih.gov/6603656/, doi: 10.1126/SCIENCE.6603656.

Graf W, Baker R. Neuronal adaptation accompanying metamorphosis in the flatfish. Journal of Neurobiology. 1990; 21(7):1136–1152. doi: 10.1002/neu.480210716.

Graf W, Spencer R, Baker H, Baker R. Excitatory and inhibitory vestibular pathways to the extraocular motor nu-clei in goldfish. Journal of neurophysiology. 1997; 77(5):2765–2779. https://pubmed.ncbi.nlm.nih.gov/9163391/, doi: 10.1152/JN.1997.77.5.2765.

Graf W, Spencer R, Baker H, Baker R. Vestibuloocular reflex of the adult flatfish. III. A species-specific reciprocal pattern of excitation and inhibition. Journal of Neurophysiology. 2001; 86(3):1376–1388. doi: 10.1152/jn.2001.86.3.1376.

Gu Z. Complex heatmap visualization. iMeta. 2022 sep; 1(3):e43. https://onlinelibrary.wiley.com/doi/full/10.1002/imt2.43 https://onlinelibrary.wiley.com/doi/abs/10.1002/imt2.43 https://onlinelibrary.wiley.com/doi/10.1002/imt2.43, doi: 10.1002/IMT2.43.

Gu Z, Eils R, Schlesner M. Complex heatmaps reveal patterns and correlations in multidimensional genomic data. Bioin-formatics. 2016 sep; 32(18):2847–2849. https://dx.doi.org/10.1093/bioinformatics/btw313, doi: 10.1093/BIOINFORMAT-ICS/BTW313.

Guerrero-Peña L, Suarez-Bregua P, Gil-Gálvez A, Naranjo S, Méndez-Martínez L, Tur R, García-Fernández P, Tena JJ, Rotl-lant J, GEO; 2023. https://identifiers.org/geo/GSE215396.

Guerrero-Peña L, Suarez-Bregua P, Sánchez-Ruiloba L, Méndez-Martínez L, García-Fernández P, Tur R, Tena JJ, Rotllant J. Unraveling the transcriptomic landscape of eye migration and visual adaptations during flatfish metamorphosis. Communications Biology 2024 7:1. 2024 mar; 7(1):1–9. https://www.nature.com/articles/s42003-024-05951-x, doi: 10.1038/s42003-024-05951-x.

Hahne F, Ivanek R. Visualizing genomic data using Gviz and bioconductor. Methods in Molecular Biology. 2016; 1418:335–351. https://link.springer.com/protocol/10.1007/978-1-4939-3578-9_16, doi: 10.1007/978-1-4939-3578-9_16/FIGURES/11.

Hauser FE, Chang BS. Insights into visual pigment adaptation and diversity from model ecological and evolutionary systems. Current Opinion in Genetics Development. 2017 dec; 47:110–120. doi: 10.1016/J.GDE.2017.09.005.

Hoke KL, Evans BI, Fernald RD. Remodeling of the Cone Photoreceptor Mosaic during Metamorphosis of Flounder (Pseu-dopleuronectes americanus). Brain, Behavior and Evolution. 2006; 68(4):241–254. https://www.karger.com/Article/FullText/94705, doi: 10.1159/000094705.

Illingworth RS, Gruenewald-Schneider U, Webb S, Kerr ARW, James KD, Turner DJ, Smith C, Harrison DJ, Andrews R, Bird AP. Orphan CpG islands identify numerous conserved promoters in the mammalian genome. PLoS genetics. 2010 sep; 6(9). https://pubmed.ncbi.nlm.nih.gov/20885785/, doi: 10.1371/JOURNAL.PGEN.1001134.

de Jesus EG, Hirano T, Inui Y. Flounder metamorphosis: its regulation by various hormones. Fish Physiology and Bio-chemistry. 1993; 11(1–6):323–328. doi: 10.1007/BF00004581.

Jia Q, Nie H, Yu P, Xie B, Wang C, Yang F, Wei G, Ni T. HNRNPA1-mediated 3 UTR length changes of HN1 contributes to cancer- and senescence-associated phenotypes. Aging (Albany NY). 2019 jul; 11(13):4407. /pmc/articles/PMC6660030//pmc/articles/PMC6660030/?report=abstract https://www.ncbi.nlm.nih.gov/pmc/articles/PMC6660030/, doi: 10.18632/AGING.102060.

Jones PL, Veenstra GJC, Wade PA, Vermaak D, Kass SU, Landsberger N, Strouboulis J, Wolffe AP. Methylated DNA and MeCP2 recruit histone deacetylase to repress transcription. Nature genetics. 1998; 19(2):187–191. https://pubmed.ncbi.nlm.nih.gov/9620779/, doi: 10.1038/561.

Kölsch Y, Hahn J, Sappington A, Stemmer M, Fernandes AM, Helmbrecht TO, Lele S, Butrus S, Laurell E, Arnold-Ammer I, Shekhar K, Sanes JR, Baier H. Molecular classification of zebrafish retinal ganglion cells links genes to cell types to behavior. Neuron. 2021 feb; 109(4):645–662.e9. http://www.cell.com/article/S0896627320309624/fulltext http://www.cell.com/article/S0896627320309624/abstract https://www.cell.com/neuron/abstract/S0896-6273(20)30962-4, doi: 10.1016/J.NEURON.2020.12.003/ATTACHMENT/679D51D5-D507-4B82-AF86-2DD2B4C9BEB5/MMC4.XLSX.

Kosmaoglou M, Kanuga N, Aguilà M, Garriga P, Cheetham ME. A dual role for EDEM1 in the processing of rod opsin. Journal of Cell Science. 2009 dec; 122(24):4465. pmc/articles/PMC2787460//pmc/articles/PMC2787460/?report=abstract https://www.ncbi.nlm.nih.gov/pmc/articles/PMC2787460/, doi: 10.1242/JCS.055228.

Krueger F, James F, Ewels P, Afyounian E, Weinstein M, Schuster-Boeckler B, Hulselmans G, FelixKrueger/TrimGalore: v0.6.10. Zenodo; 2023. doi: 10.5281/zenodo.5127898.

Kulczyńska K, Siatecka M. A regulatory function of long non-coding RNAs in red blood cell development. Acta biochimica Polonica. 2016; 63(4):675–680. https://pubmed.ncbi.nlm.nih.gov/27851834/, doi: 10.18388/ABP.2016_1351.

Kupkova K, Mosquera JV, Smith JP, Stolarczyk M, Danehy TL, Lawson JT, Xue B, Stubbs JT, LeRoy N, Sheffield NC. Ge-nomicDistributions: fast analysis of genomic intervals with Bioconductor. BMC Genomics. 2022 dec; 23(1):1–6. https://bmcgenomics.biomedcentral.com/articles/10.1186/s12864-022-08467-y, doi: 10.1186/S12864-022-08467-Y/FIGURES/2.

Kyono Y, Raj S, Sifuentes CJ, Buisine N, Sachs L, Denver RJ. DNA methylation dynamics underlie metamorphic gene regulation programs in Xenopus tadpole brain. Developmental Biology. 2020; 462(2):180–196. https://doi.org/10.1016/j.ydbio.2020.03.013, doi: 10.1016/j.ydbio.2020.03.013.

Kyono Y, Sachs LM, Bilesimo P, Wen L, Denver RJ. Developmental and thyroid hormone regulation of the DNA methyltrans-ferase 3a gene in xenopus tadpoles. Endocrinology. 2016 dec; 157(12):4961–4972. /pmc/articles/PMC5133355//pmc/articles/PMC5133355/?report=abstract https://www.ncbi.nlm.nih.gov/pmc/articles/PMC5133355/, doi: 10.1210/EN.2016-1465/SUPPL_FILE/EN-16-1465.PDF.

Lamb TD. Evolution of phototransduction, vertebrate photoreceptors and retina. Progress in Retinal and Eye Research. 2013 sep; 36:52–119. doi: 10.1016/J.PRETEYERES.2013.06.001.

Lawrence M, Huber W, Pagès H, Aboyoun P, Carlson M, Gentleman R, Morgan MT, Carey VJ. Software for Computing and Annotating Genomic Ranges. PLOS Computational Biology. 2013; 9(8):e1003118. https://journals.plos.org/ploscompbiol/article?id=10.1371/journal.pcbi.1003118, doi: 10.1371/JOURNAL.PCBI.1003118.

Liu J, Reggiani JDS, Laboulaye MA, Pandey S, Chen B, Rubenstein JLR, Krishnaswamy A, Sanes JR. Tbr1 instructs laminar patterning of retinal ganglion cell dendrites. Nature Neuroscience. 2018; 21(5):659–670. https://doi.org/10.1038/s41593-018-0127-z, doi: 10.1038/s41593-018-0127-z.

Love MI, Huber W, Anders S. Moderated estimation of fold change and dispersion for RNA-seq data with DESeq2. Genome Biology. 2014 ec; 15(12):550. doi: 10.1186/s13059-014-0550-8.

Lu CF, Zhou YN, Zhang J, Su S, Liu Y, Peng GH, Zang W, Cao J. The role of epigenetic methylation/demethylation in the regulation of retinal photoreceptors. Frontiers in Cell and Developmental Biology. 2023; 11:1149132. /pmc/articles/PMC10251769//pmc/articles/PMC10251769/?report=abstract https://www.ncbi.nlm.nih.gov/pmc/articles/PMC10251769/, doi: 10.3389/FCELL.2023.1149132.

Mader MM, Cameron DA. Photoreceptor differentiation during retinal development, growth, and regeneration in a metamorphic vertebrate. Journal of Neuroscience. 2004; 24(50):11463–11472. doi: 10.1523/JNEUROSCI.3343-04.2004.

Mao CA, Kiyama T, Pan P, Furuta Y, Hadjantonakis AK, Klein WH. Eomesodermin, a target gene of Pou4f2, is required for retinal ganglion cell and optic nerve development in the mouse. Development (Cambridge, England). 2008 jan; 135(2):271. /pmc/articles/PMC2893890//pmc/articles/PMC2893890/?report=abstract https://www.ncbi.nlm.nih.gov/pmc/articles/PMC2893890/, doi: 10.1242/DEV.009688.

Maunakea AK, Nagarajan RP, Bilenky M, Ballinger TJ, Dsouza C, Fouse SD, Johnson BE, Hong C, Nielsen C, Zhao Y, Turecki G, Delaney A, Varhol R, Thiessen N, Shchors K, Heine VM, Rowitch DH, Xing X, Fiore C, Schillebeeckx M, et al. Conserved role of intragenic DNA methylation in regulating alternative promoters. Nature 2010 466:7303. 2010 jul; 466(7303):253–257. https://www.nature.com/articles/nature09165, doi: 10.1038/nature09165.

McKiernan PJ, Greene CM. The Biology of Long Non-Coding RNA. MicroRNAs and Other Non-Coding RNAs in Inflam-mation. 2015 jan; p. 21–42. https://link.springer.com/chapter/10.1007/978-3-319-13689-9_2, doi: 10.1007/978-3-319-13689-9_2.

Medina M, Rep J, Ward R, Rio P, Lemire M. The primary visual system of flatfish: an evolutionary perspective. Techniques. 1993; p. 167–191.

Meehan RR, Stancheva I. DNA methylation and control of gene expression in vertebrate development. Essays in bio-chemistry. 2001; 37:59–70. https://pubmed.ncbi.nlm.nih.gov/11758457/, doi: 10.1042/BSE0370059.

Merbs SL, Khan MA, Hackler L, Oliver VF, Wan J, Qian J, Zack DJ. Cell-specific DNA methylation patterns of retina-specific genes. PloS one. 2012 mar; 7(3). https://pubmed.ncbi.nlm.nih.gov/22403679/, doi: 10.1371/JOURNAL.PONE.0032602.

Muñoz-Mérida A, Viguera E, Claros MG, Trelles O, Pérez-Pulido AJ. Sma3s: A three-step modular annotator for large sequence datasets. DNA Research. 2014; 21(4):341–353. doi: 10.1093/dnares/dsu001.

Murray M. Axonal transport in the asymmetric optic axons of flatfish. Experimental Neurology. 1974 mar; 42(3):636–646. https://linkinghub.elsevier.com/retrieve/pii/0014488674900855, doi: 10.1016/0014-4886(74)90085-5.

Neant I, Deisig N, Scerbo P, Leclerc C, Moreau M. The RNA-binding protein Xp54nrb isolated from a Ca2+-dependent screen is expressed in neural structures during Xenopus laevis development. The International journal of develop-mental biology. 2011; 55(10–12):923–931. https://pubmed.ncbi.nlm.nih.gov/22252489/, doi: 10.1387/IJDB.103253IN.

Okano M, Bell DW, Haber DA, Li E. DNA methyltransferases Dnmt3a and Dnmt3b are essential for de novo methyla-tion and mammalian development. Cell. 1999 oct; 99(3):247–257. https://pubmed.ncbi.nlm.nih.gov/10555141/, doi: 10.1016/S0092-8674(00)81656-6.

Rao RC, Hennig AK, Malik MTA, Chen DF, Chen S. Epigenetic regulation of retinal development and disease. Journal of Ocular Biology, Diseases, and Informatics. 2011 sep; 4(3):121. /pmc/articles/PMC3382290//pmc/articles/PMC3382290/?report=abstract https://www.ncbi.nlm.nih.gov/pmc/articles/PMC3382290/, doi: 10.1007/S12177-012-9083-0.

Robles E, Laurell E, Baier H. The retinal projectome reveals brain-area-specific visual representations generated by ganglion cell diversity. Current Biology. 2014 sep; 24(18):2085–2096. http://www.cell.com/article/S0960982214009816/fulltext http://www.cell.com/article/S0960982214009816/abstract https://www.cell.com/current-biology/abstract/S0960-9822(14)00981-6, doi: 10.1016/j.cub.2014.07.080.

Rottach A, Leonhardt H, Spada F. DNA methylation-mediated epigenetic control. Journal of cellular biochemistry. 2009 sep; 108(1):43–51. https://pubmed.ncbi.nlm.nih.gov/19565567/, doi: 10.1002/JCB.22253.

Sara Frau, Iñigo Novales Flamarique, Patrick W Keeley, Benjamin E Reese JAMC. Straying from the flatfish retinal plan: cone photoreceptor patterning in the common sole (Solea solea) and the Senegalese sole (Solea senegalensis). Physiology behavior. 2017; 176(3):139–148. doi: 10.1053/j.gastro.2016.08.014.CagY.

Schreiber AM. Asymmetric craniofacial remodeling and liberalized behavior in larval flatfish. Journal of Experimental Biology. 2006; 209(4):610–621. doi: 10.1242/jeb.02056.

Shao C, Bao B, Xie Z, Chen X, Li B, Jia X, Yao Q, Ortí G, Li W, Li X, Hamre K, Xu J, Wang L, Chen F, Tian Y, Schreiber AM, Wang N, Wei F, Zhang J, Dong Z, et al. The genome and transcriptome of Japanese flounder provide insights into flatfish asymmetry. Nature Genetics. 2017; 49(1):119–124. doi: 10.1038/ng.3732.

Suarez-Bregua P, Pérez-Figueroa A, Hernández-Urcera J, Morán P, Rotllant J. Temperature-independent genome-wide DNA methylation profile in turbot post-embryonic development. Journal of Thermal Biology. 2020; 88(November 2019):1–7. doi: 10.1016/j.jtherbio.2019.102483.

Tai WL, Ashok A, Cho KS, Guan TP, Cho J, An A, Chen D. Regulation of retinal ganglion cell axon growth and optic nerve regeneration by DNA methyltransferase. Investigative Ophthalmology Visual Science. 2023 jun; 64(8):2840–2840.

Tata JR. Amphibian metamorphosis as a model for studying the developmental actions of thyroid hormone. Cell Research. 1998; 8(4):259–272. doi: 10.1038/cr.1998.26.

Tian H, He Y, Xue Y, Gao YQ. Expression regulation of genes is linked to their CpG density distributions around transcrip-tion start sites. Life Science Alliance. 2022 sep; 5(9). https://www.life-science-alliance.org/content/5/9/e202101302 https://www.life-science-alliance.org/content/5/9/e202101302.abstract, doi: 10.26508/LSA.202101302.

Vance KW, Ponting CP. Transcriptional regulatory functions of nuclear long noncoding RNAs. Trends in Genetics. 2014; 30(8):348. /pmc/articles/PMC4115187//pmc/articles/PMC4115187/?report=abstract https://www.ncbi.nlm.nih.gov/pmc/articles/PMC4115187/, doi: 10.1016/J.TIG.2014.06.001.

Villegas EDM, Dans MJD, Castillo CPA, Alvarez RA. Development of the eye in the turbot Psetta maxima (Teleosti) from hatching through metamorphosis. Journal of Morphology. 1997; 233(1):31–42. doi: 10.1002/(SICI)1097-4687(199707)233:1<31::AID-JMOR3>3.0.CO;2-R.

Wang Y, Zhou L, Wu L, Song C, Ma X, Xu S, Du T, Li X, Li J. Evolutionary ecology of the visual opsin gene sequence and its expression in turbot (Scophthalmus maximus). BMC Ecology and Evolution. 2021; 21(1):1–12. https://doi.org/10.1186/s12862-021-01837-2, doi: 10.1186/s12862-021-01837-2.

Wu H, Xu T, Feng H, Chen L, Li B, Yao B, Qin Z, Jin P, Conneely KN. Detection of differentially methylated regions from whole-genome bisulfite sequencing data without replicates. Nucleic Acids Research. 2015 ec; 43(21):e141. /pmc/articles/PMC4666378//pmc/articles/PMC4666378/?report=abstract https://www.ncbi.nlm.nih.gov/pmc/articles/PMC4666378/, doi: 10.1093/NAR/GKV715.

Xi Y, Li W. BSMAP: Whole genome bisulfite sequence MAPping program. BMC Bioinformatics. 2009 jul; 10(1):1–9. https://bmcbioinformatics.biomedcentral.com/articles/10.1186/1471-2105-10-232, doi: 10.1186/1471-2105-10-232/COMMENTS.

Yu G, Wang LG, Han Y, He QY. ClusterProfiler: An R package for comparing biological themes among gene clusters. OMICS A Journal of Integrative Biology. 2012 may; 16(5):284–287. doi: 10.1089/omi.2011.0118.

